# Thymic stromal lymphopoietin contributes to endometriotic lesion proliferation and disease-associated inflammation

**DOI:** 10.1101/2024.04.18.590132

**Authors:** Stanimira Aleksieva, Harshavardhan Lingegowda, Danielle J Sisnett, Alison McCallion, Katherine B Zutautas, Timothy Childs, Bruce Lessey, Chandrakant Tayade

**Author notes:** Corresponding author: Dr. Chandrakant Tayade^1^, DVM, PhD, Department of Biomedical and Molecular Sciences Queen’s University, Kingston, ON, Canada, K7L 3N6.

## Abstract

Endometriosis is a chronic disorder in which endometrial-like tissue presents outside the uterus. Patients with endometriosis have been shown to exhibit aberrant immune responses within the lesion microenvironment and in circulation which contribute to the development of endometriosis. Thymic stromal lymphopoietin (TSLP) is an alarmin involved in cell proliferation and the induction of Th2 inflammation in various diseases, such as asthma, atopic dermatitis, pancreatic and breast cancer. Recent studies have detected TSLP within endometriotic lesions and shown that its concentrations are elevated in the peritoneal fluid of patients compared to controls. However, its role in disease pathophysiology remains unclear. Here, we compared TSLP expression in endometriotic lesions to matched patient endometrium and control endometrium samples. We also assessed its effect on the proliferation and apoptosis of human endometriosis-representative cell lines, as well as on lesion development and inflammation in a mouse model of the disease. We demonstrated that TSLP expression was elevated in the stroma of patient endometriotic lesions compared to control endometrial samples. In cell lines, TSLP treatment reduced the apoptosis of endometrial stromal cells and promoted the proliferation of THP-I cells. In mice induced with endometriosis, TSLP treatment induced a Th2 immune response within the lesion microenvironment, and led to TSLP receptor modulation in macrophages, dendritic cells, and CD4+ T cells. Furthermore, treatment increased murine endometriotic lesion proliferation. Overall, these results suggest that TSLP modulates the endometriotic lesion microenvironment and promotes a Th2 immune response that could support lesion development.

**KEY POINTS:**

- TSLP is overexpressed in the stroma of human endometriotic lesions;
- TSLP promotes the survival of human endometrial stromal and THP-I cell lines;
- TSLP induces Th2 inflammation and lesion proliferation.

## INTRODUCTION

Endometriosis is a chronic disorder that affects an estimated 200 million women worldwide (1). In endometriosis, tissue similar in composition to the endometrium, termed endometriotic lesions, grows outside of the uterus. This often results in chronic pelvic pain independent of menstrual cyclicity (2). Despite the prevalence of the disorder and its impact on women’s health, the cause remains elusive. We and others have provided evidence that an altered immune response could hinder lesion clearance and facilitate lesion development through promotion of tissue adhesion, invasion, proliferation, and angiogenesis (2, 3). Some examples of altered responses include increased alternative macrophage polarization and enhanced T helper (Th) 2 immune responses in the lesion microenvironment (4–8).

One important mediator of Th2 immune responses is thymic stromal lymphopoietin (TSLP) (9, 10). TSLP is an alarmin that can be secreted from multiple sources, such as stromal, epithelial, and innate immune cells in response to shear stress or inflammation from stimuli such as IL-1β, IL-4, and IL-13 (9, 11, 12). TSLP interacts with the TSLP receptor (TSLPR) and the interleukin IL-7 receptor α (IL-7Rα) chain. Dimerization activates JAK 1, 2, STAT 1, 3, and 5, which are implicated in the proliferation of immune and stromal cells, such as CD4+ T cells, and activates Th2 responses that involve the production of interleukin (IL)-4, −5, −9, −13 (9). These end products further serve as stimuli for TSLP secretion, which enables the formation of a positive feedback loop that accentuates Th2 inflammation (13). Th2 inflammation is important to maintain homeostasis in response to Th1 inflammation, which involves IL-1β, IFN-γ, and TNF-α. However, in excess, Th2 responses have been linked to the initiation and exacerbation of numerous chronic conditions, such as asthma, atopic dermatitis, pancreatic and breast cancer (9, 10, 14–16).

In endometriosis, TSLP mRNA and protein have been detected within lesions, and shown to be elevated in the peritoneal fluid and serum of patient samples compared to control samples (12, 17). Furthermore, various stimuli present within the lesion microenvironment, such as IL-1β, IL-4, and 17-β-estradiol, can induce TSLP secretion from endometrial stromal cells (12, 18). Nevertheless, the contribution of TSLP to lesion development and the immune response in endometriosis remains unclear. This is critical to address as Th2 responses promote lesion adherence, invasion, proliferation, and survival (14).

Here, we address the involvement of TSLP in endometriosis pathophysiology using patient samples, endometriosis-representative cell lines, and a syngeneic C57BL/6 mouse model of endometriosis. Our results reveal that TSLP is over-expressed in patient lesion stroma, contributes to the survival of endometriotic stromal cells, and promotes lesion proliferation and Th2 responses in mice induced with endometriosis.

## MATERIALS/METHODS

### Ethics statement

Ethics for the use of human samples was approved by the Health Sciences Research Ethics Board at Kingston Health Sciences Centre and Queen’s University (Kingston, Canada). Samples were collected from patients with and without endometriosis with informed consent before collection and research application. Animal studies and protocols were approved by the Queen’s University Animal Care Committee (Kingston, Canada).

### Immunohistochemistry for TSLP on an endometrioma tissue microarray

To establish whether TSLP protein expression varies between endometriosis patients and controls, a tissue microarray (TMA) with matched endometriosis patient eutopic and ectopic tissues (n = 19) and non-endometriosis control endometrium (n = 15) was assessed. All patient lesions were endometriomas – lesions that occur on the ovaries. Full sample characteristics are described in Sisnett et al. (19). The TMA was deparaffinized with xylene and re-hydrated with CitriSolv and graded concentrations of alcohol. Antigen retrieval was performed with EDTA buffer for 30 min. The slide was stained with a polyclonal anti-TSLP Ab (ab47943, Abcam, UK; 1:100) and the BOND Polymer Refine Detection Kit (DS9800, Leica Biosystems, USA; contains diaminobenzidine chromogen and hematoxylin counterstain) using a Leica Bond RX automated slide stainer (Leica Biosystems, USA). The slide was scanned with an Olympus VS120 Virtual Slide Microscope (Olympus, USA) at 20X magnification. A custom tissue classifier and area quantification algorithm was created with HALO AI Software (Indica Labs, USA) to assess TSLP-positive stroma and epithelia as a fraction of total stromal and epithelial area, respectively.

#### Human cell lines

### Human endometriotic epithelial cells (12Zs)

Immortalized human endometriotic epithelial cells (12Z; Prof. Anna Starzinski-Powitz, Goethe University, Germany) were cultured in DMEM/F-12 (11330; ThermoFisher Scientific) supplemented with 10% fetal bovine serum (FBS; 97068-085, VWR), 1% penicillin/streptomycin solution (15140122, ThermoFisher), and 1X sodium pyruvate (11360070, ThermoFisher).

### Human endometrial stromal cells (hESCs)

Immortalized human endometrial stromal cells (hESC; T0533, Applied Biological Materials, Canada) were cultured in Prigrow IV growth media (TM004, Applied Biological Materials, Canada) supplemented with 10% charcoal-stripped FBS (12676029, ThermoFisher Scientific, Canada), 1% L-glutamine (A2916801, ThermoFisher Scientific, Canada), and 1% penicillin/streptomycin solution.

### Human monocytic leukemia (THP-I) cells

Human monocytic leukemia cells (THP-I; TIB-202TM, American Type Culture Collection, USA) were cultured in standard media comprised of RPMI-1640 medium (11875085, ThermoFisher Scientific, Canada) supplemented with 10% FBS (FBS; 97068-085, VWR). All cell lines were maintained within a standard incubator at 37°C with 5% CO_2_.

#### Proliferation and apoptosis assays in cell lines treated with recombinant human TSLP

12Z, hESC, and THP-I cells were seeded at 5 X 10^3^ cells/well in 96-well plates with 100 μL of their respective media. After 24 h of rest, cells were treated in triplicates with vehicle control (PBS; 10% v/v) or recombinant human TSLP (rhTSLP; 1398-TS-010/CF, R&D Systems, USA) (1, 5, 10, 50, 100 ng/mL) for 24 h. To establish whether TSLP treatment impacts the cell lines’ proliferation, wells were treated with 10 μL of WST-I reagent (501594401, Millipore Sigma, Canada) for 2 h at 37°C with 5% CO_2_. Absorbance, proportional to the amount of metabolically-active cells in culture, was quantified with the SpectraMaxiD3 microplate reader (Molecular Devices, USA) at a wavelength reading of 440 nm. To establish whether treatment impacts the apoptosis of endometriosis-relevant cell lines, 12Z, hESC, and THP-I cells were treated with 100 μL of Caspase-Glo 3/7 reagent (G8091, Promega, Canada) for 2 h at room temperature. Luminescence, proportional to caspase 3/7-mediated apoptosis, was quantified with the SpectraMaxiD3 microplate reader. All results were calculated after subtracting background absorbance or luminescence from a cell-free media control.

#### Treatment of THP-I cells with TSLP and flow cytometry

To understand whether TSLP treatment influences the polarization of THP-I cells, cells were seeded at 1 X 10^6^ cells in 6-well plates. Cells were cultured in standard media supplemented with 20 nM phorbol 12-myristate 13-acetate (PMA; P1585, Sigma Aldrich, USA) for 48 h, then rested in standard media for another 48 h to induce differentiation into adherent, activated macrophage-like cells. Differentiated cells were treated in triplicates with vehicle control (PBS; 10% v/v) or various concentrations of rhTSLP (10, 50, 100 ng/mL) for 24 h. After treatment, adherent cells were detached from plates with versine solution (15040066, ThermoFisher Scientific, Canada) as per manufacturer’s protocol. Cells were centrifuged at 500*g* for 5 min at 4°C to pellet, re-suspended in PBS supplemented with 2% FACS buffer (PBS with 2% FBS), and aliquoted for flow cytometric analysis (5 X 10^5^ cells/sample).

#### Harvest of mouse bone marrow progenitors, differentiation into bone-marrow derived macrophages, treatment with TSLP, and flow cytometry

To understand whether TSLP treatment also affects the polarization of mouse bone marrow-derived macrophages (BMDMs), bone marrow progenitors were harvested from C57BL/6 mouse (Charles River Laboratories, USA) femurs as previously detailed (20). Cells were seeded at 5 X 10^5^ cells in 6-well plates and cultured in RPMI-1640 (11875085, ThermoFisher Scientific, Canada) supplemented with 2% HEPES (H0887, Sigma Aldrich, USA), 10% FBS (FBS; 97068-085, VWR), 1% 100X non-essential amino acids (11140050, ThermoFisher Scientific, Canada), 5% 200 nM L-glutamine (25030081, ThermoFisher Scientific, Canada), 5% 100X penicillin/streptomycin solution (15140122, ThermoFisher Scientific, Canada), 0.0025% 200 µg/mL M-CSF (576402, BioLegend, USA) to induce differentiation into macrophages. Cells were treated in triplicates with vehicle control (PBS) or various concentrations of recombinant mouse TSLP (rmTSLP; 555-TS-010/CF, R&D, USA; 1, 10, 50 ng/mL) for 7 d at 37°C with 5% CO_2_.

Post-treatment, supernatant was aliquoted and stored at −80°C. Cells were detached from plates with ice-cold versine solution (15040066, ThermoFisher Scientific, Canada) as per manufacturer’s protocol, centrifuged at 500*g* for 5 min at 4°C, re-suspended in 2% FACS, and aliquoted into tubes for flow cytometric analysis (5 X 10^5^ cells/sample).

#### Syngeneic mouse model of endometriosis

Six-to-eight-week-old female immunocompetent C57BL/6 mice were housed in conventional housing. To prepare for endometriosis induction, 4 mm^3^ endometrial tissue fragments were obtained from the uterine horns of donor mice (n = 3) with an epidermal punch tool, and 100 µL of blood was collected from the submandibular vein of recipient mice (n = 9) and into EDTA-coated tubes (395974, BD Biosciences, USA). Blood was centrifuged at 3000 RCF for 15 min at 4°C, and plasma was aliquoted for multiplex cytokine analysis. To induce endometriosis, recipient mice were anesthetized with 3.5% vaporized isoflurane, a small abdominal incision was made, two donor uterine fragments were adhered to the peritoneal wall with VetBond tissue adhesive (1469SB, 3M, USA), and the abdomen was closed with sutures and staples. On 7 d postoperative, staples were removed, and from 8d to 18d postoperative, mice were injected i.p. with vehicle control (n = 5; PBS; 100 µL) or rmTSLP (n = 5; 55-TS-010/CF, R&D Systems, USA; 2 µg or 0.01 mg/kg) every other day for a total of six times (Fig. S3A).

On 20 d postoperative, 100 µL of blood was collected from the submandibular vein of mice, then mice were euthanized. Peritoneal fluid was collected *via* a peritoneal lavage with 4 mL ice-cold PBS, lesions were excised and placed in 4% paraformaldehyde, and spleen was harvested and placed in RPMI-1640 (11875085, ThermoFisher Scientific, Canada) supplemented with 10% FBS. Blood was process as outlined above. Peritoneal fluid was centrifuged at 500*g* for 5 min at 4°C, then fluid was stored at −80 °C for multiplex cytokine analysis, and cells were resuspended in 2% FACS for flow cytometric analysis. Spleens were mechanically digested using 70 µm strainers and resuspended in ACK Lysing Buffer (A1049201, ThermoFisher Scientific), as per manufacturer’s instruction. Then, cells were centrifuged at 500*g* for 5 min at 4°C and cells were re-suspended in 2% FACS for flow cytometric analysis. To assess how TSLP treatment influences the local and systemic immune response, two aliquots of peritoneal fluid cells and splenocytes were made to conduct flow cytometric analysis (5 X 10^5^ cells/sample). Endometriotic lesions were stored in paraformaldehyde for 24h, transferred to ethanol, and embedded in paraffin for immunohistochemical analysis.

#### Flow cytometric analysis

Processed cells were incubated with human or mouse TruStain FcX (101320, 422302, BioLegend, USA), based on cell source, for 5 min at 4°C to prevent non-specific binding. Cells were stained with the extracellular abs and incubated for 30 min at 4°C prior (Table I). Then, cells were permeabilized with the FOXP3 Fixation and Permeabilization Kit (00552300, ThermoFisher Scientific, Canada) as per manufacturer’s protocol, stained with the intracellular Ab and incubated for 30 min at room temperature (Table I). The CytoFLEX S flow cytometer (Beckman Coulter, USA) was used to analyze stained cells. Data was analyzed with FlowJo version 10 Software (FlowJo, USA).

**Table I.**
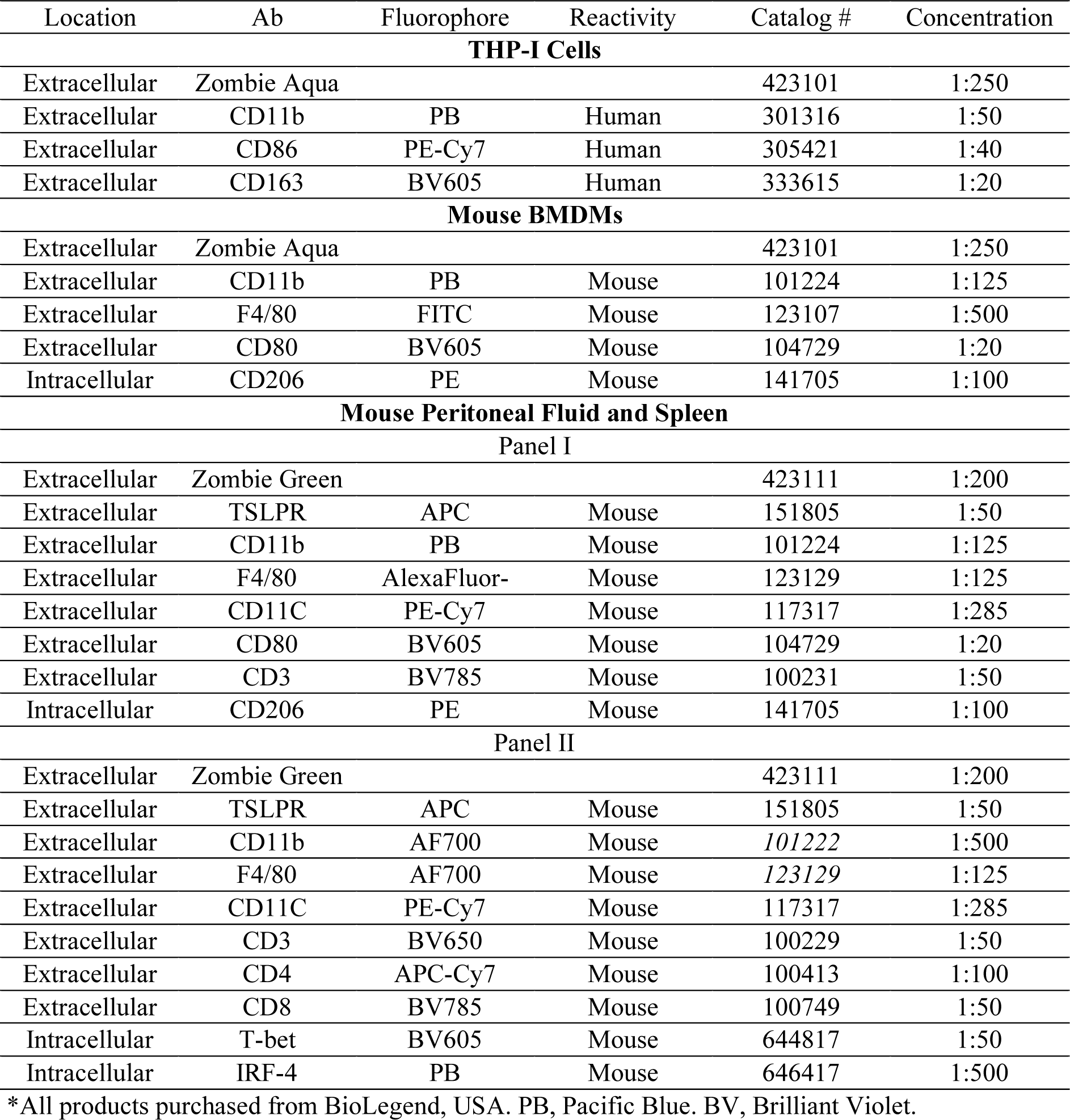
List of extracellular and intracellular Abs used for flow cytometric analysis.

#### Multiplex cytokine analysis to detect prominent cytokines and chemokines in mouse BMDM cell culture and peritoneal fluid

To detect cytokines released in response to rmTSLP treatment, 100 µL samples of mouse BMDM cell culture supernatant, peritoneal fluid and plasma underwent multiplex cytokine array to measure eotaxin, G-CSF, GM-CSF, IFN-*γ*, IL-1α, IL-1β, IL-2, IL-3, IL-4, IL-5, IL-6, IL-7, IL-9, IL-10, IL-12 (p40), IL-12 (p70), IL-13, IL-15, IL-17, IP-10, KC, LIF, LIX, MCP-1, M-CSF, MIG, MIP-1α, MIP-1β, MIP-2, RANTES, TNFα, and VEGF (MD32, Eve Technologies, CA).

#### Immunohistochemistry for Ki67, CD31, and TSLPR expression within mouse endometriotic lesions

Embedded mouse endometriotic lesions were sectioned at 5 μm thickness. Sections were deparaffinized with xylene and re-hydrated with CitriSolv and graded concentrations of alcohol. Antigen retrieval was performed with citrate buffer for 30 min. Slides were stained with anti-Ki67 (ab15580, Abcam, USA; 1:2000), anti-CD31 (77699S, New England Biolabs, USA; 1:600), and anti-TSLPR (BS-4652R, ThermoFisher Scientific, Canada, 1:200) abs and the BOND Polymer Refine Detection Kit (DS9800, Leica Biosystems, USA) using the Leica Bond RX automated slide stainer (Leica Biosystems, USA). Slides were scanned with the Olympus VS120 Virtual Slide Microscope (Olympus, USA) at 20X magnification. Images were assessed with HALO AI Software (Indica Labs, USA). Two custom area quantification algorithms were created with HALO AI Software (Indica Labs, USA) to detect CD31- and TSLPR-positive area as a fraction of total tissue area. One custom quantification algorithm was also generated to detect Ki67-positive cells as a fraction of total cells in tissue.

### Statistical analysis

Statistical analysis was conducted with GraphPad Prism9 software (GraphPad Software Inc., USA). Data are represented as mean ± SD. An unpaired, non-parametric Student t test was conducted to compare between two treatment conditions, whereas an ordinary, one-way ANOVA with Tukey post-hoc test was used to compare between three or more treatment conditions. A *P* value below 0.05 was considered significant.

## RESULTS

### TSLP is elevated in stromal tissue of endometrioma lesions

To understand the spatial localization and expression of TSLP, immunohistochemistry was conducted on a TMA constructed with matched patient endometrial and endometrioma (i.e., lesions) tissue (n = 19) and control endometrial tissue (n = 15) as detailed previously (3, 19). TSLP was detected across all patient and control tissues (Fig. 1A-C). TSLP expression was significantly elevated in patient endometriomas as compared to control endometrial samples (Fig. 1D). TSLP expression did not differ in epithelial compartments between patients and controls (Fig. 1E). While TSLP has been previously localized within stromal and epithelial cells, this is the first quantification of altered TSLP protein expression levels between patient endometrioma lesions and control endometrial samples.

**Figure 1:**
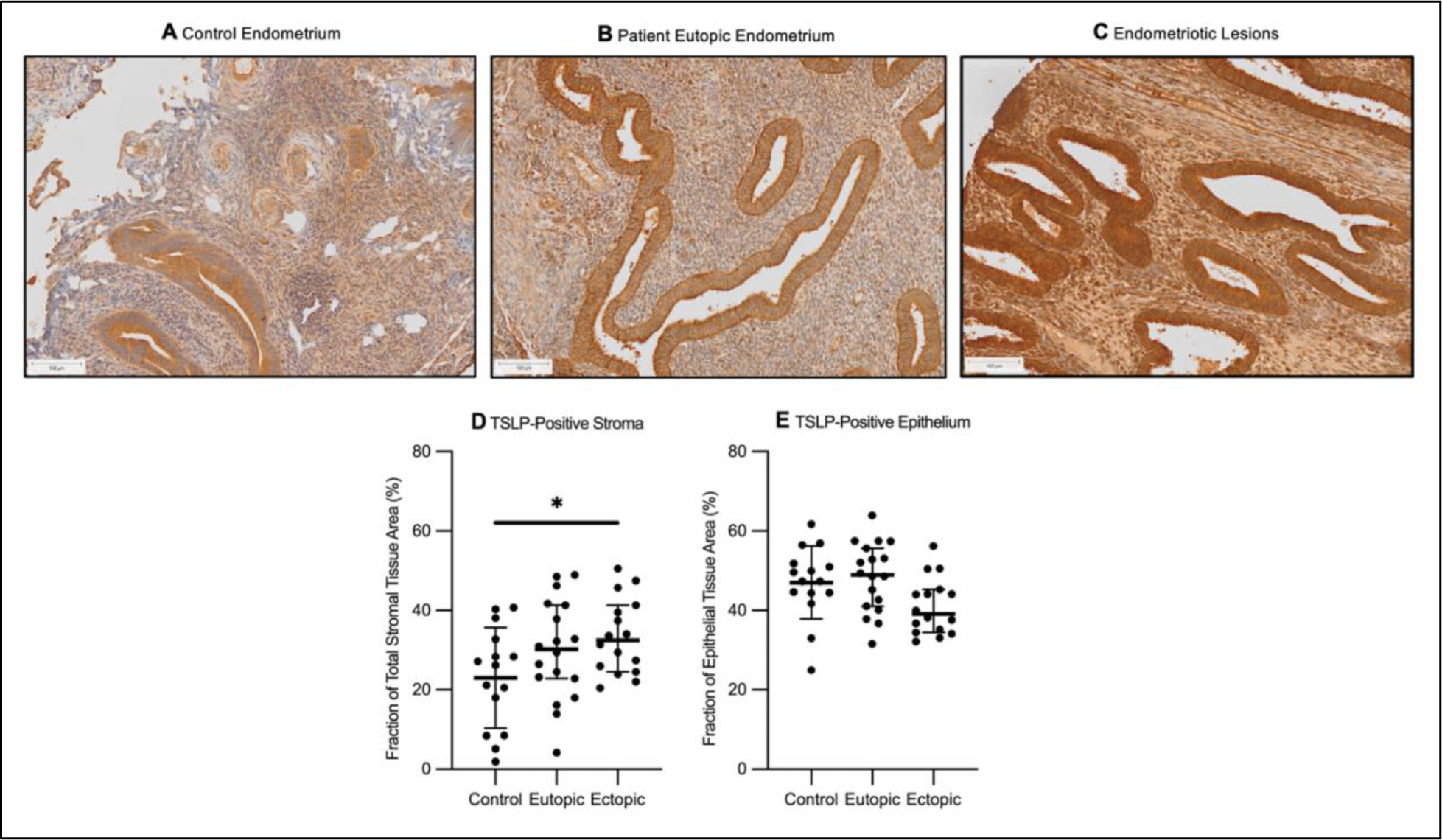
TSLP protein is significantly increased in patient endometriomas compared to healthy endometrial samples. A TMA with (**A**) control endometrial tissue (n = 15), and matched (**B**) patient endometrial (n = 19) and (**C**) endometrioma tissue was stained with anti-TSLP antibody. TSLP-positive (**D**) stromal and (**E**) epithelial area was calculated as a fraction of total stromal and epithelial area, respectively, using HALO AI (Indica Labs). Magnification provided at 8X. Scale bars 100 μm. A one-way ANOVA with Tukey post-hoc was used to assess statistical significance. \**P*<0.05.

### TSLP treatment influences the survival of endometriotic-representative cell lines in vitro

To understand the effects of rhTSLP treatment on the proliferation and apoptosis of endometriosis-representative cell lines, WST-I and caspase-3/7 assays were conducted on 12Zs, hESCs, and THP-I cells. TSLP treatment did not influence the proliferation (Fig. 2A) or apoptosis (Fig. 2B) of 12Z cells. Furthermore, TSLP did not affect the proliferation of hESCs (Fig. 2C) but did decrease their apoptosis (Fig. 2D) compared to PBS control. TSLP treatment also increased the proliferation (Fig. 2E) of THP-I cells compared to PBS control.

**Figure 2:**
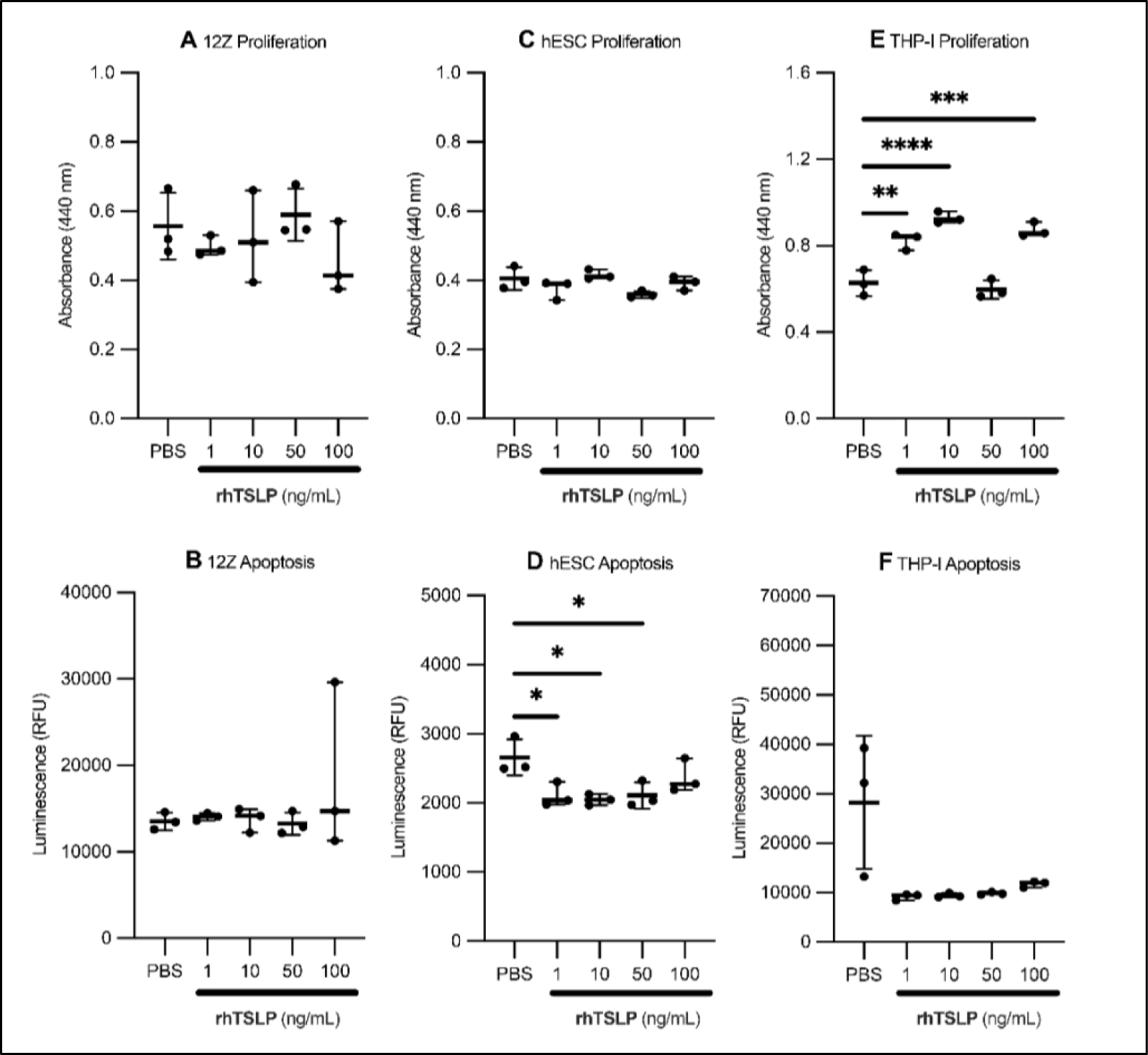
TSLP treatment alters the proliferation and apoptosis of human endometriosis-representative cell lines. (**A, B**) 12Zs, (**C, D**) hESCs, and (**E, F**) THP-I cells were treated with vehicle control (PBS) or various concentrations of rhTSLP (1, 10, 50 ng/mL) for 24h. (**A, C, E**) WST-I proliferation and (**B, D, F)** caspase-3/7 apoptosis assays were performed. A one-way ANOVA with Tukey post-hoc was used to assess statistical significance. \**P* <0.05, \*\**P* <0.01, \*\*\**P* <0.001, and \*\*\*\**P* <0.0001.

### TSLP treatment modulates surface marker expression on THP-I cells and mouse BMDMs

To evaluate the effects of TSLP treatment on the polarization of the human THP-I cell line and mouse BMDMs cells were treated with vehicle control or various concentrations of recombinant TSLP (Fig. S1, S2). In THP-I cells, TSLP treatment did not impact the fraction of CD11b+ cells (Fig. 3A) but increased the ratio of CD163+ to CD68+ cells (Fig. 3B). In mouse BMDMs, TSLP treatment did not alter the fraction of CD11b+F4/80+ (Fig. 3C) but led to a noticeable, but not significant increase in the ratio of CD206+CD80-to CD80+CD206-cells (Fig. 3D). The increased expression of alternative (CD163+ and CD206+) to classical (CD86+ and CD80+) macrophage activation markers suggests that TSLP likely enhances type 2 immune responses.

**Figure 3:**
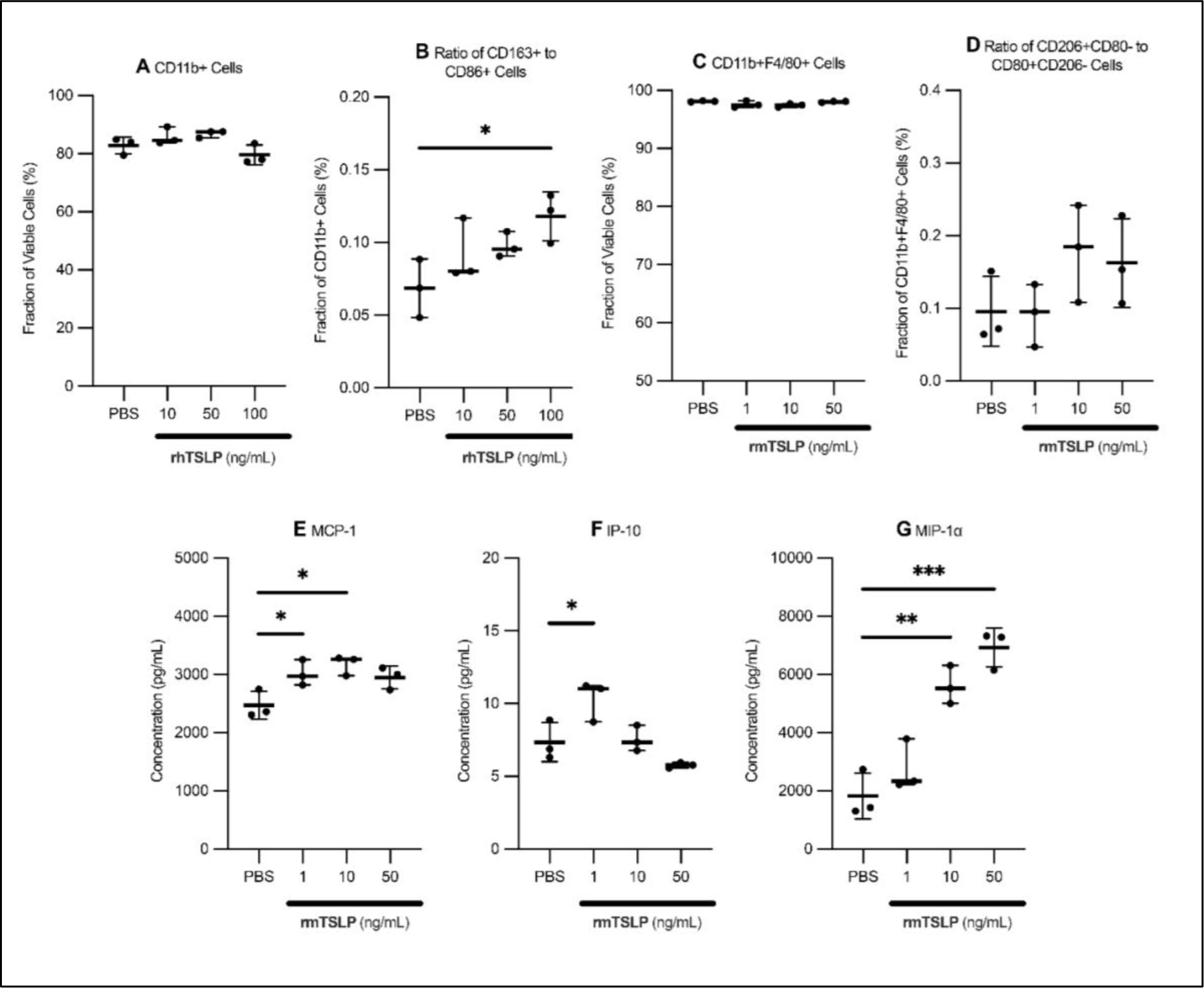
TSLP treatment influences the polarization of THP-I cells and mouse BMDMs. THP-I cells were treated with vehicle control (PBS) or various concentrations of rhTSLP (1, 10, 50 ng/mL) for 1 d, and mouse BMDMs were treated with vehicle control (PBS) or various concentrations of rmTSLP (1, 10, 50 ng/mL) for 7 d. **(A)** Flow cytometry was used to detect the expression of (**B**) CD11b, (**C**) CD163, and CD86 on THP-I cells, as well as (**D**) CD11b, F4/80, (**E**) CD80, and CD206 on BMDMs. Cell supernatant was collected from BMDMs and analyzed to detect the expression of several inflammation- and angiogenesis-related cytokines and chemokines, including (**F**) MCP-1, (**G**) IP-10, and (**H**) MIP-1α. A one-way ANOVA with Tukey post-hoc was conducted to assess statistical significance. \**P* <0.05, \*\**P* < 0.01, and \*\*\**P* <0.001.

To further understand the effects of TSLP treatment on macrophage activation, we assessed the effects of TSLP treatment on the secretion of chemokines and cytokines from mouse BMDMs. TSLP treatment significantly increased levels of monocyte chemoattractant protein-1 (MCP-1) (Fig. 3E), IFN-gamma-inducible protein 10 (IP-10) (Figure 3F), and macrophage inflammatory protein-1 alpha (MIP-1α) (Fig. 3G). These changes suggest TSLP could stimulate immune cell recruitment *through* macrophages.

### TSLP treatment in a syngeneic mouse model of endometriosis alters local and systemic immune cell populations

To understand the effects of TSLP on the local and systemic immune microenvironment and on lesion development, we induced endometriosis in C57BL/6 mice and treated with vehicle control (PBS) or rmTSLP (2 µg/dose, or 0.01 mg/kg) six times over fourteen days (Fig. S3A). To capture changes in local and systemic inflammation, peritoneal fluid cells and splenocytes were collected and analyzed with flow cytometry (Fig. S3B-L).

In peritoneal fluid, rmTSLP treatment did not alter the fraction (Fig. 4A) or polarization (data not shown) of CD11b+F4/80+ cells but reduced TSLPR expression (Fig. 4B). Furthermore, it did not affect the fraction of CD11C+ cells (Fig. 4C) or TSLPR (Fig. 4D) expression but increased their interferon regulatory factor 4 (IRF-4) expression (Fig. 4E). rmTSLP treatment also increased the proportion of CD4+ T cells (Fig. 4F), TSLPR (Fig. 4G) expression, and IRF-4 to T-bet ratio (Fig. 4H). No differences in CD8+ T cell numbers (Fig. 4I) or TSLPR expression (Fig. 4J) were observed. These changes suggest that TSLP predominantly promotes Th2 cell activation.

**Figure 4:**
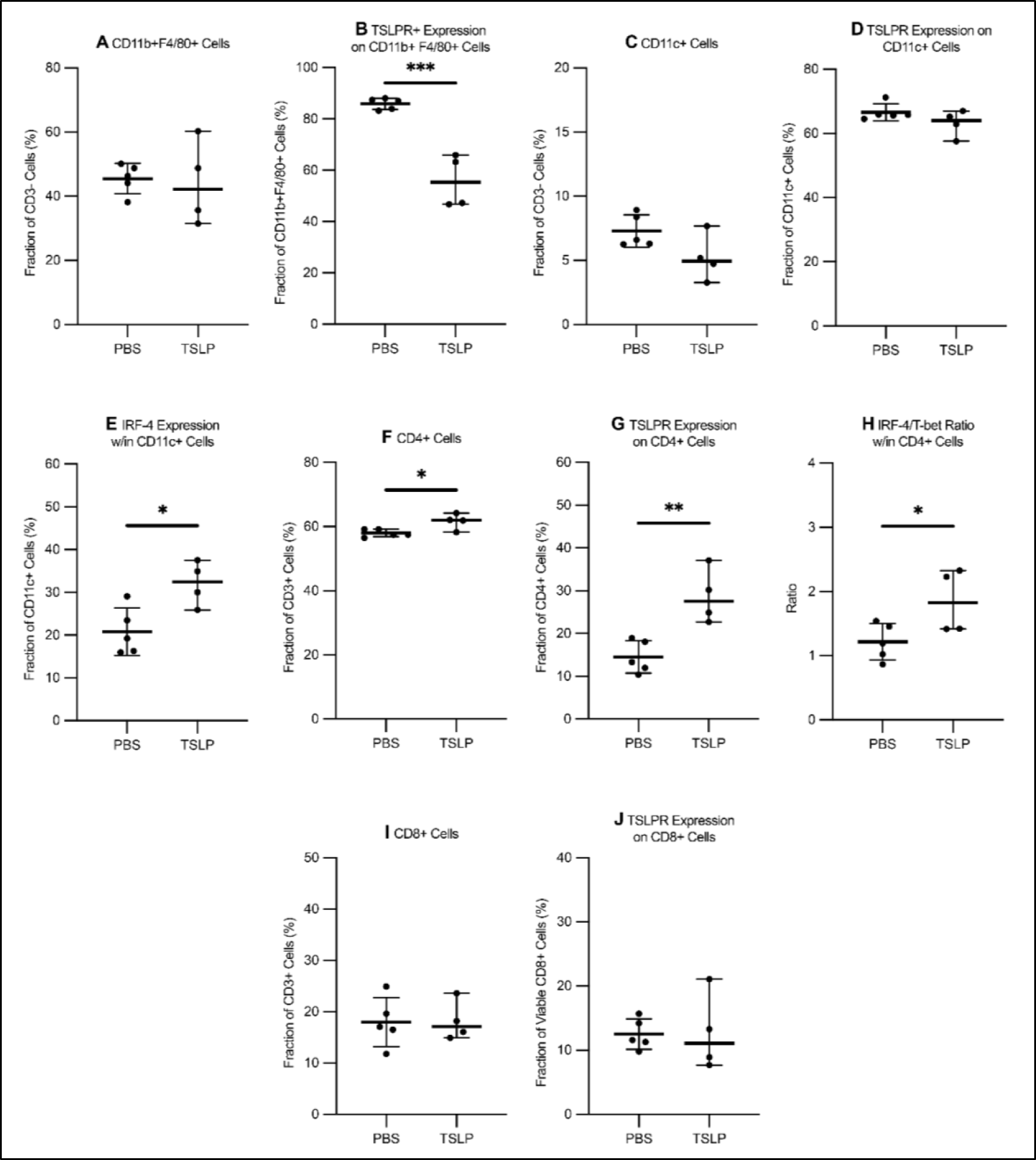
TSLP treatment influences the presence and activation of immune cells in the peritoneal fluid of mice induced with endometriosis. Mice were treated with i.p. PBS (n = 5) or rmTSLP (2 μg or 0.01 mg/kg; n = 4) every other day for 14 days. (**A-E**) Myeloid and (**F-J**) lymphoid cells, as well as their (**B, D, G, J**) TSLPR expression, (**E**) IRF4 expression, and (**F, J**) IRF-4/T-bet ratio was measured. An unpaired student’s T-test was used to assess statistical significance. * *P*<0.05, \*\**P* < 0.01, and \*\*\**P* <0.001

In the spleen, rmTSLP treatment did not affect the fraction of CD11C+ cells (Fig. 5A), nor their TSLPR (Fig. 5B) and IRF-4 (Fig. 5C) expression. TSLP treatment also did not impact the number of CD4+ cells (Fig. 5D) but reduced TSLPR (Fig. 5E) expression and increased their IRF-4/T-bet ratio (Fig. 5F). Furthermore, treatment reduced the presence of CD8+ T cells (Fig. 5G), and their TSLPR (Fig. 5H) expression. These results suggest that TSLP alters the systemic immune microenvironment in a manner distinct from the peritoneal microenvironment.

**Figure 5:**
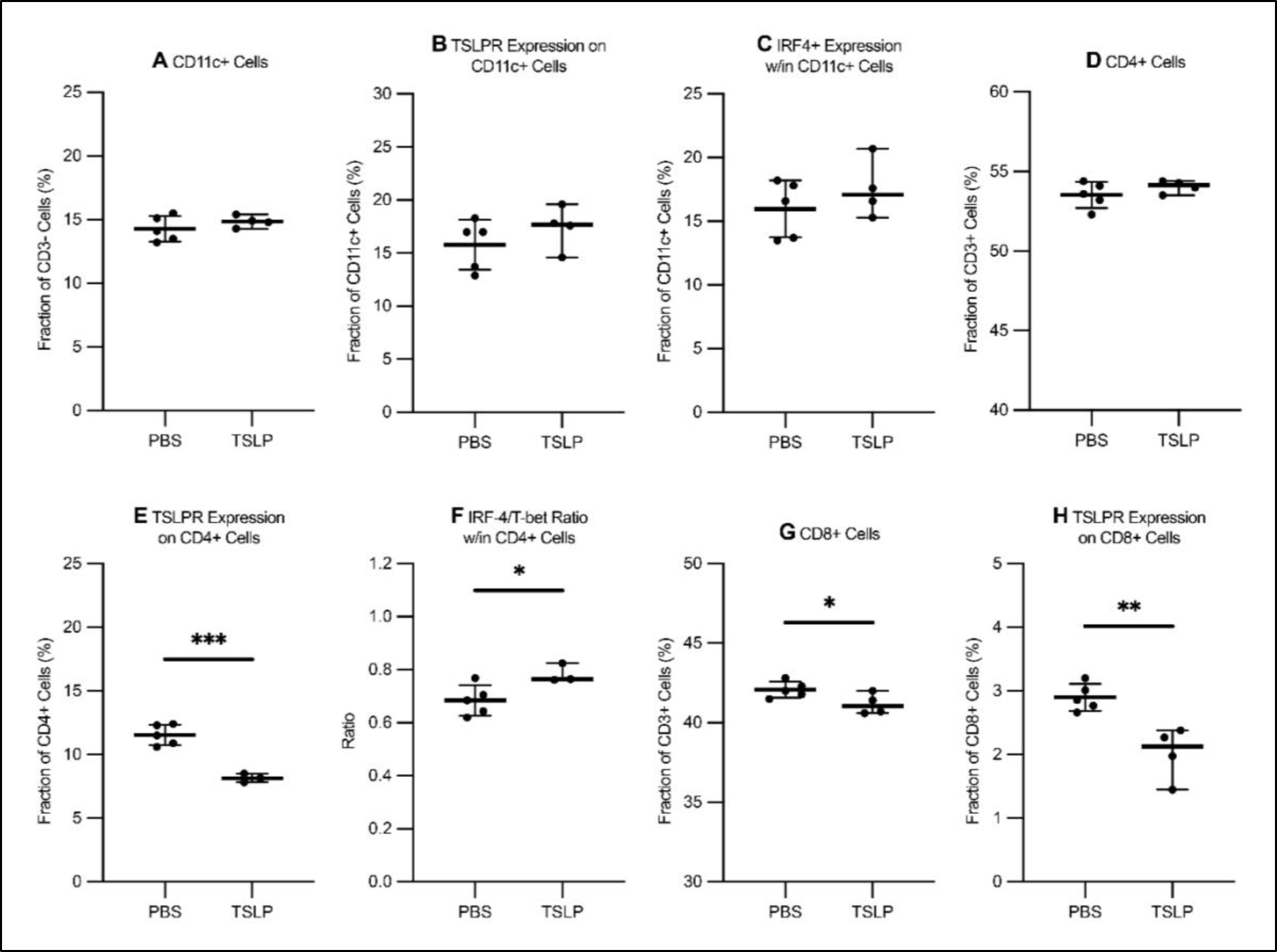
TSLP treatment influences the presence and activation of immune cells in circulation. Mice were treated with i.p. PBS (n = 5) or rmTSLP (2 μg or 0.01 mg/kg; n = 4) every other day for 14 days. (**A-C**) Myeloid and (**D-H**) lymphoid cells, and their (**B, E, H**) TSLPR expression, IRF-4 expression, and (**F, J**) IRF-4/T-BET ratio was measured. An unpaired student’s T-test was used to assess statistical significance. \**P* <0.05, \*\**P* <0.01, \*\*\**P* <0.001.

To further capture changes in local and systemic inflammation, peritoneal fluid supernatant and plasma were analyzed with a multiplex cytokine array for predominant pro- and anti-inflammatory cytokines, chemokines, and growth factors. In peritoneal fluid, rmTSLP treatment reduced the concentration of IL-6 (Fig. 6A) and IFN-γ (Fig. 6B) but did not affect the levels of IL-10 (Fig. 6C) and IL-13 (Fig. 6D). No difference in these mediators was observed in plasma (data not shown).

**Figure 6:**
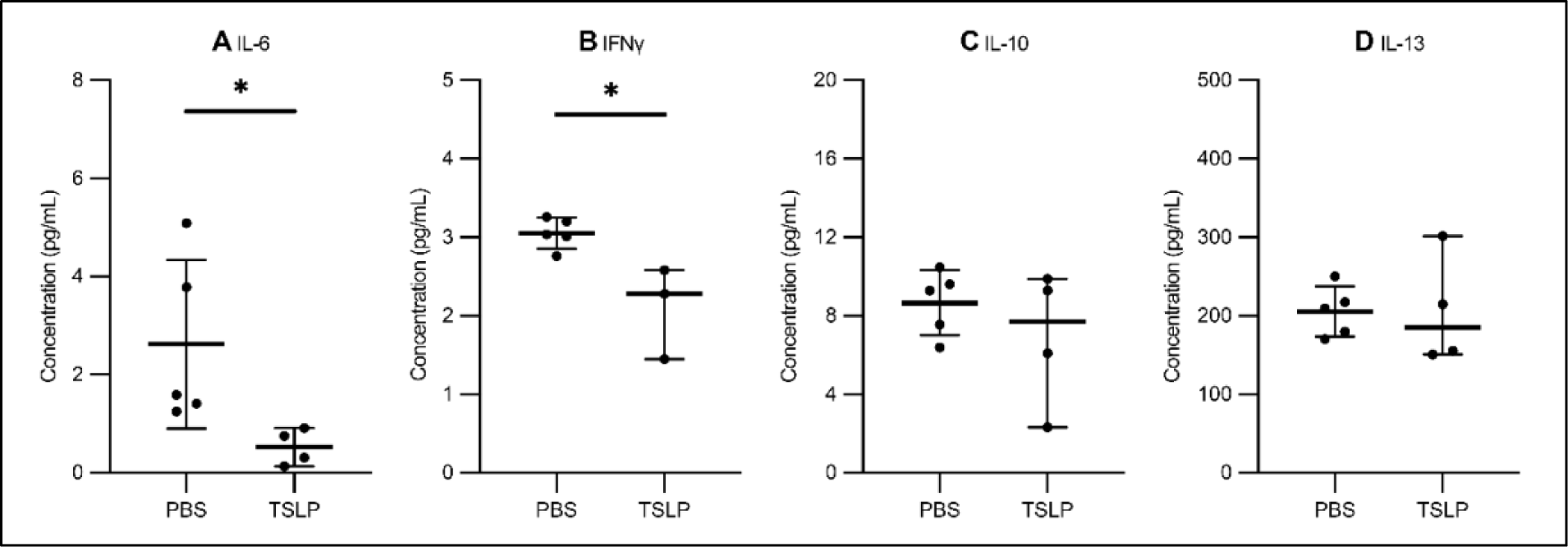
TSLP treatment reduces the concentrations of prevalent pro-inflammatory cytokines in mouse peritoneal fluid. Mice were treated with i.p. PBS (n = 5) or rmTSLP (2 μg or 0.01 mg/kg; n =4) every other day for 14 days. Peritoneal fluid was collected to detect predominant inflammatory cytokines and chemokines. Key (**A, B)** pro- and **(C, D**) anti-inflammatory cytokines shown. An unpaired student’s T-test was conducted to assess statistical significance. \**P* <0.05.

### TSLP treatment in a syngeneic mouse model of endometriosis promotes lesion proliferation

TSLP modules TSLPR expression in immune cells, proliferation in endometrial stromal cells, and angiogenesis in cervical cancer (21). To see whether TSLP treatment also influences lesion TSLPR expression, proliferation, and angiogenesis, lesions were processed and stained with anti-TSLPR, anti-Ki67, and anti-CD31 Ab, respectively. TSLP treatment did not alter TSLPR (Fig. 7A) or CD31 (Fig. 7C) levels, however Ki67 expression increased (Fig. 7B).

**Figure 7:**
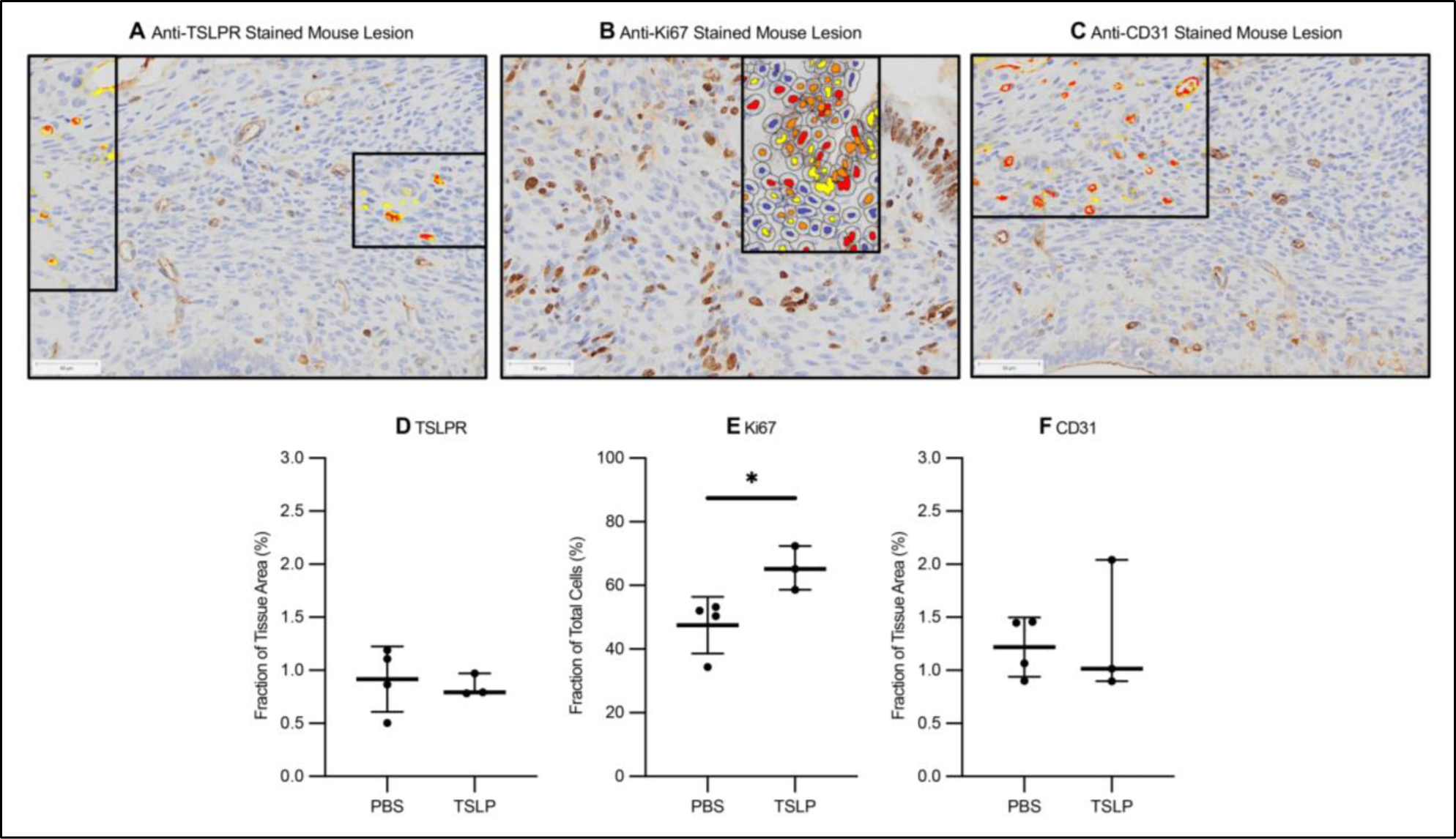
TSLP treatment significantly increases murine lesion proliferation. Endometriotic lesions from mice treated with i.p. PBS (n = 5) or rmTSLP (2 μg or 0.01 mg/kg; n = 4) were embedded in paraffin and stained with anti-**(A)** TSLPR, **(B)** anti-Ki67, and **(C)** anti-CD31 Abs. TSLPR-, **(E)** Ki67-, and **(F)** CD31-positive tissue was calculated with HALO AI (Indica Labs). Magnification provided at 20X. Scale bars 50 μm. An unpaired student’s T-test was used to assess statistical significance. \**P* <0.05.

## DISCUSSION

Current literature shows that elevated TSLP potentiates the development of chronic auto-immune conditions and inflammatory conditions, including asthma, atopic dermatitis, rheumatoid arthritis, several cancers, such as breast and pancreatic (9, 10). Here, we provide evidence for its relevance in endometriosis by evaluating its expression in patient as well as control samples and examining its influence on lesion proliferation and disease-associated inflammation.

In the past, TSLP was shown to be expressed within endometrioma lesions, and to be elevated in the peritoneal fluid and serum of endometriosis patients (12). Furthermore, elevated TSLP mRNA expression was reported in patient endometriotic lesions, as compared to matched eutopic endometrium and control endometrial samples (17). However, a comparison of TSLP protein expression between patient lesions and control endometrium had not been conducted. Here, we identified elevated TSLP expression in the stroma of endometriotic lesions as compared to that of control endometrium. The increased expression corresponds to previous work which localized TSLP within the stroma and demonstrated that the human endometrial cell line can upregulate TSLP secretion in response to various inflammatory and hormonal stimuli, such as IL-1β, IL-4, and 17β-estradiol (12, 22). Furthermore, we observed no differences in TSLP protein expression in the epithelium of endometriotic lesions. While previous research localized and reported more intense TSLP stain near the epithelium as compared to the stroma of endometrioma lesions, our automated quantification demonstrated no differences in expression TSLP-positive area in the epithelium of patients versus controls (12). Overall, these results provide new, quantitative insight into the unique microenvironment of endometriotic lesions.

Previous research has shown that TSLP treatment can promote the proliferation of human endometrial stromal cells, bronchial epithelial cells, and CD4+ T cells (22–24). Furthermore, in animal models of asthma and chronic skin inflammation, it enhances the recruitment of immune cells, *such as* macrophages (25, 26). In addition, based on evidence from cell cultures and a mouse model of asthma, TSLP is postulated to interact with macrophages and enhance their alternative polarization (25, 27). Here, we demonstrated that TSLP reduces hESC apoptosis and increases THP-I proliferation. In addition, TSLP treatment promoted the differentiation of human THP-Is and mouse BMDMs towards an alternatively-activated phenotype, and stimulated mouse BMDMs to secrete numerous chemokines, such as MCP-1 and MIP-1α. Macrophages are an abundant immune cell population within the lesion microenvironment that can assume a broad spectrum of activation and functional states based on local microenvironment cues. On opposite ends of the spectrum lie classically and alternatively subsets, which can be identified based on numerous extracellular and intracellular markers (28). Whereas the former pro-inflammatory stimuli, such as IL-1β, IFN-γ, and have been linked to lesion clearance, the latter secrete anti-inflammatory stimuli, such as IL-4 and −13, and promote lesion development (28, 29). In our work, the reduced apoptosis of our stromal cell line corresponds to previous work which has shown that TSLP treatment can reduce stromal cell apoptosis and prolong their viability (18, 22, 30). The polarization of THP-Is and BMDMs is also in line with reports of TSLP-mediated THP-I maturation and alternative murine alveolar macrophage activation (25, 31). The observed release of chemokines from mouse BMDMs further suggests that TSLP could mediate immune cell recruitment through macrophage activation.

To understand the effects of TSLP on the local and systemic immune microenvironment, as well as on lesion development, we used our established immunocompetent mouse model of endometriosis treated with rmTSLP. In peritoneal fluid, TSLP treatment reduced TSLPR expression but did not affect macrophage polarization. This differs from a TSLP-treated mouse model of asthma, where TSLP over-expression in lung tissue increased TSLPR expression and promoted alternative polarization, with the polarization response dependent upon on the increased TSLPR expression (25). While there are various factors which could account for the different response, such as differences in TSLP treatment conditions, tissue-specific differences in response, it is also possible that TSLPR downregulation in our model is an adaptive response to maintain immune homeostasis in response to TSLP challenge. TSLP treatment also did not influence dendritic cell proportions but increased their IRF-4 expression. Dendritic cells are known to express TSLPR, and TSLP treatment is well-known induce the expression of numerous co-stimulatory molecules, including OX40L, CD80, and CD86 (10, 32, 33). As IRF-4 expression within dendritic cells is linked to their secretion of IL-10 and −13 this increase exemplifies another means through which TSLP could be implicated in Th2 polarization (10, 34, 35). Th2 responses involve the production of anti-inflammatory mediators, such as IL-4 and IL-10 (14). These are considered antagonistic to Th1 responses which involve pro-inflammatory mediators, such as IL-2, IL-6, and IFN-γ. In endometriosis, the two responses coexist, albeit one could predominate based on numerous factors, such as the site of endometriosis, duration of disease, the hormonal milieu, and individual differences in immune function. Nevertheless, it is accepted that in advanced endometriosis, there is a shift towards a Th2 phenotype that progress disease through the promotion of lesion proliferation, angiogenesis, inflammation, and 17β-estradiol production. In addition to the changes observed in macrophages dendritic cells, TSLP treatment increased CD4+ T cell proportions, TSLPR expression, and the intracellular ratio of the Th1 and Th2 transcription factors T-bet and IRF-4, respectively (36–41). These results correspond to previous literature which showed that TSLP can interact with CD4+ T cells to increase proliferation, survival, recruitment, and Th2 polarization (24, 26, 42–45). Furthermore, the increase in TSLPR expression corresponds with previous work that identified TSLP treatment enhances CD4+ T cell TSLPR expression, with the highest level present on Th2 cells (43). In peritoneal fluid, TSLP treatment reduced the concentration of IL-6 and IFN-γ, but not of IL-10 and IL-13. The reduction in the Th1 cytokines further suggests a shift towards Th2 responses (39, 40).

In the spleen, TSLP treatment did not influence dendritic cell proportions nor their IRF-4 expression. The fraction of CD4+ T cell was also maintained, albeit TSLP treatment reduced their TSLPR expression and increased their IRF-4 to T-bet ratio. This suggests that while TSLP may not be implicated in the modulation of dendritic cell IRF-4 expression, nor upregulate CD4+ T cell numbers and TSLPR expression, it can nevertheless promote systemic Th2 polarization. In the spleen, TSLP treatment significantly reduced CD8+ T cell proportions and TSLPR expression. Current literature suggests that TSLP interacts with the TSLPR to enhance the survival and homeostasis of naïve CD8+ T cells, but repress that of effector CD8+ T cells (46, 47). Thus, further characterization of CD8+ T cell subsets is required to understand the functional significance of our observations on endometriotic lesions.

Within our murine endometriotic lesions, we identified constitutive TSLPR expression. TSLP treatment increased lesion proliferation but not angiogenesis. The effects on proliferation correspond to our and others’ *in vitro* TSLP treatment of stromal cell lines, where TSLP enhanced proliferation (22). While TSLP has been found to promote angiogenesis in cervical cancer, here we demonstrate that these effects may not be translatable to endometriotic lesions as quantified by CD31 (21, 48).

In conclusion, we demonstrate that TSLP is upregulated in patient lesions and could contribute to lesion growth and endometriosis-associated immune dysregulation through the modulation of innate and adaptive immune responses. Nevertheless, we acknowledge the limitations of our work. First, we evaluated TSLP expression in nineteen patient endometriomas. Samples from additional endometriosis stages and phenotypes are needed to validate current results in a larger cohort of patients and assess differences in expression between subtypes. Second, while our immunocompetent mouse model of endometriosis incorporates important characteristics of endometriotic lesions, such as the presence of stroma, glandular epithelium, and cysts, it does not recapitulate natural disease establishment. This includes the process of menstruation, the hormonal and inflammatory microenvironments within or outside of the uterus that are conducive to lesion establishment, as well as the chronic nature of endometriosis. In consequence, our model does not address the extent to which TSLP-mediated effects are a cause or effect of endometriosis. This remains an important question, as previous research has shown that TSLP can stimulate Th2 responses, and vice versa. Future experiments should be implemented to understand how the inflammatory and hormonal milieu of endometriotic lesions, and how the timing of TSLP administration relative to endometriosis induction influences lesion development. Overall, our work examining the impact of TSLP in endometriosis furthers our understanding of the immunological basis of the disease and assists in the future development of effective therapies to inhibit disease progression.

