## Supplemental Figure 1 - 3. for "Thymic stromal lymphopoietin contributes to endometriotic lesion proliferation and disease-associated inflammation"

**SUPPLEMENTAL DATA**

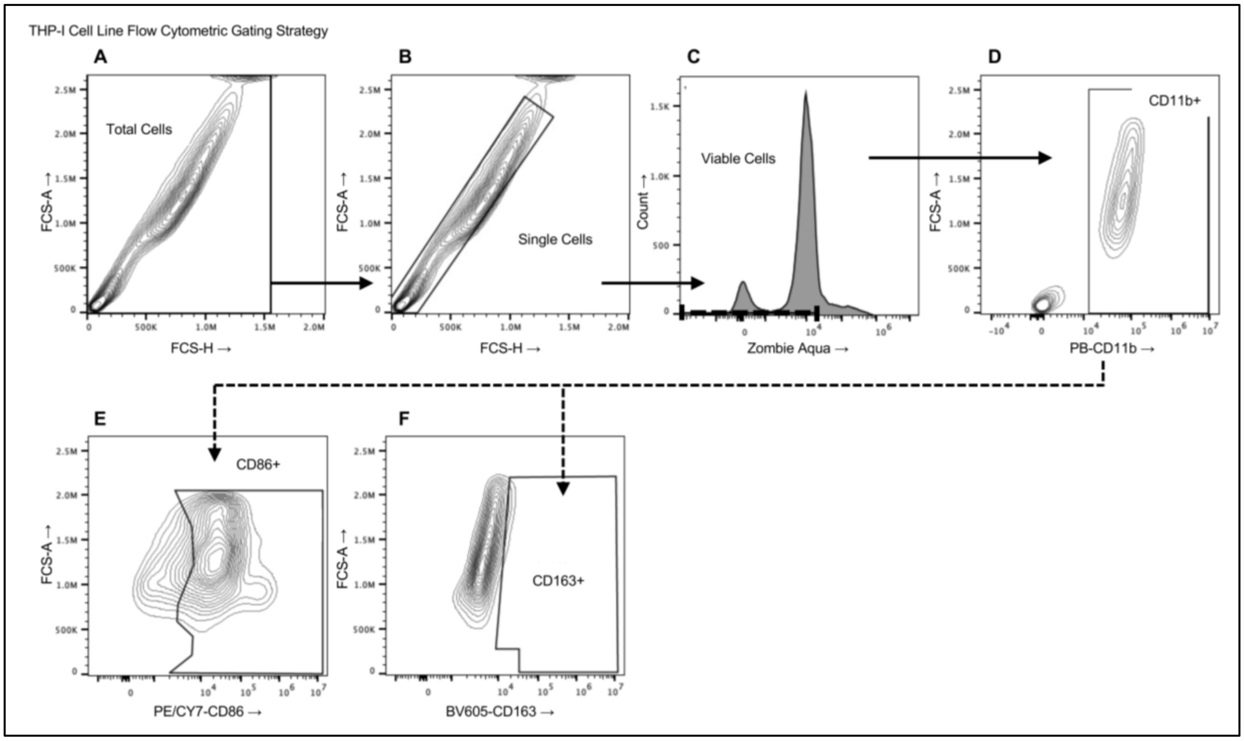


**Figure S1 |** Flow cytometric analysis of the human THP-I cell line. THP-I cells were treated with PBS or various concentrations of rhTSLP (10, 50, 100 ng/mL). Flow cytometry was used to select **(A)** total cells, **(B)** single cells, **(C)** viable cells, as well as their **(D)** CD11b, **(E)** CD86, and **(F)** CD163+ expression.

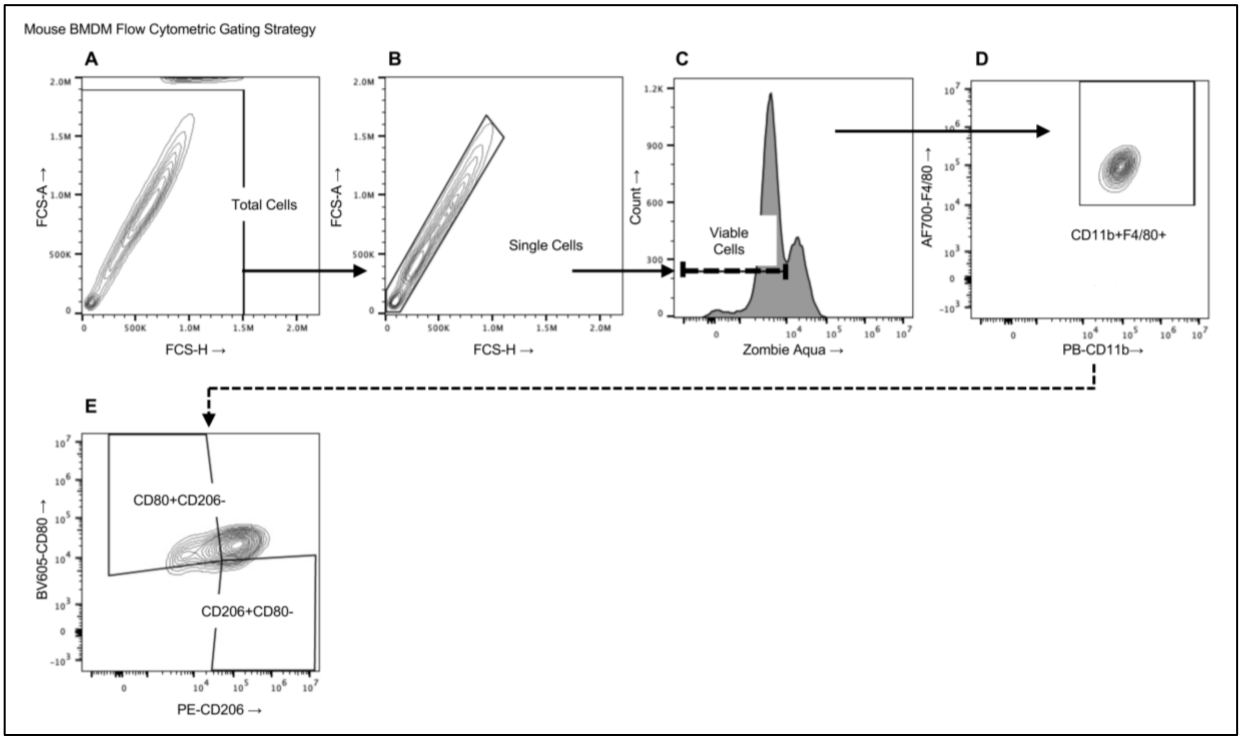


**Figure S2 |** Flow cytometric analysis of mouse BMDMs. BMDMs were treated with PBS or various concentrations of rmTSLP (1, 10, 50 ng/mL). Flow cytometry was used to select **(A)** total cells, **(B)** single cells, **(C)** viable cells, as well as their **(D)** CD11b, **(E)** CD80 and CD206 expression.

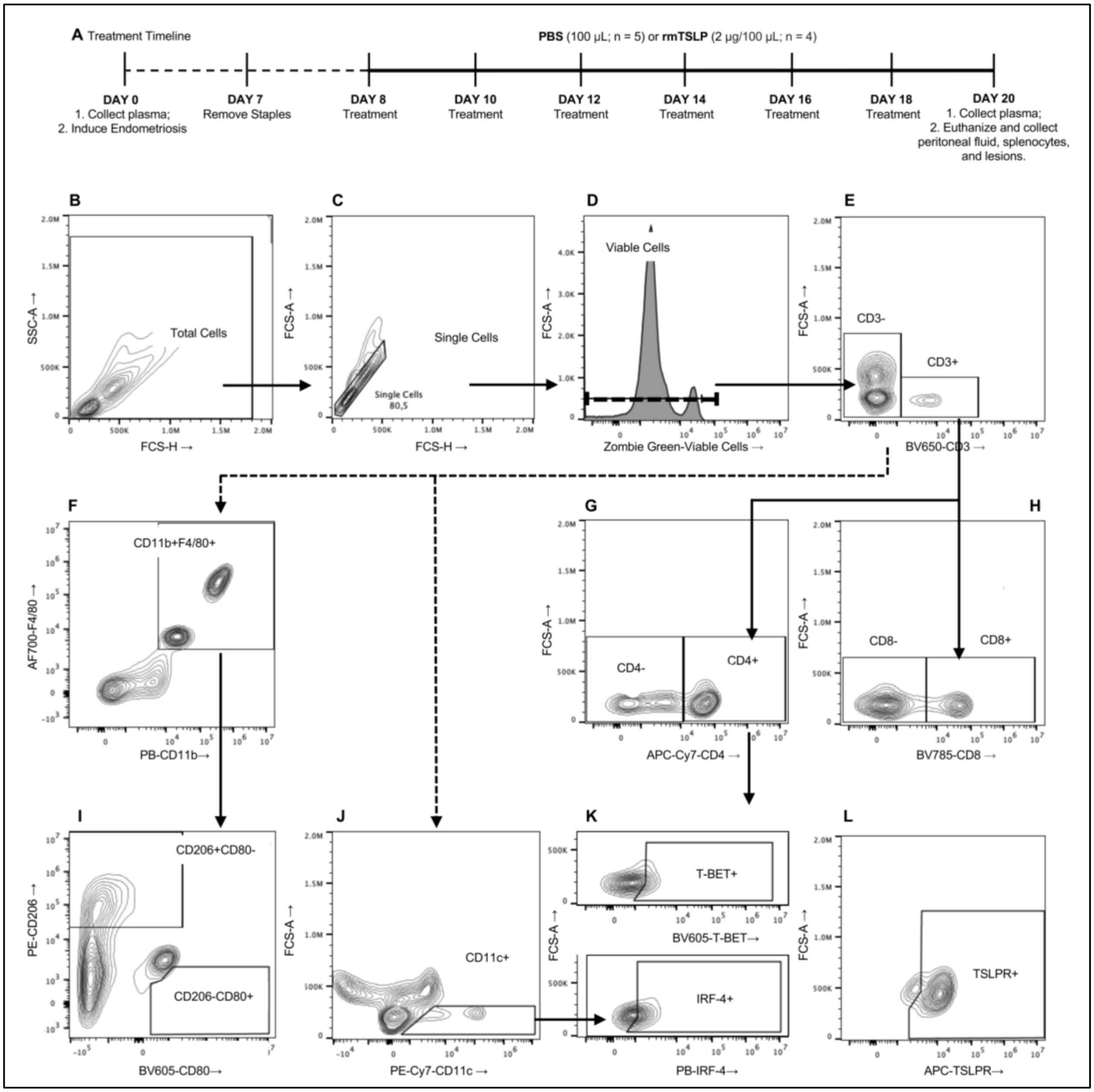


**Figure S3 |** Gating strategy for flow cytometric analysis of peritoneal fluid cells and splenocytes. **(A)** Mice were induced with endometriosis and treated with i.p PBS (n = 5) or rmTSLP (2 µg or 0.01 mg/kg; n = 4). At endpoint, peritoneal fluid and spleen were collected for flow cytometric analysis. **(B)** Total cells, **(C)** single cells, and **(D)** viable cells were selected, and the expression of (**E**) CD3, (**F**) CD11b, F4/80, (**G**) CD4, (**H**) CD8, (**I**) CD206, CD80, (**J**) CD11c, (**K**) T-bet, IRF-4, and **(L)** TSLPR was assessed.
